# A twofold strategy for the protection of therapeutic peptides by attachment to the protease-resistant and fusogenic protein, saposin C

**DOI:** 10.1101/2020.07.24.219675

**Authors:** Suzanne I. Sandin, Christopher J. Randolph, Eva de Alba

**Author notes:** Corresponding Author Eva de Alba. Department of Bioengineering. School of Engineering. University of California, Merced.

## Abstract

A great challenge of therapeutic peptides (biologics) is their short half-life. However, biologics can be protected by encapsulation in liposomes used as drug-delivery platforms. Liposomes are typically incorporated into cells by endocytic pathways, which eventually expose therapeutics to favorable proteolytic conditions. To enhance biologics protection, we report the design and characterization of a liposome-protein chimera combining the liposome fusogenic properties of peripheral-membrane protein saposin C, covalently linked to a proapoptotic peptide (the active domain of Bcl-2 protein PUMA). We show by NMR that the saposin C component of the chimera is capable of binding liposomes and that the peptide binds prosurvival Bcl-xL, thus following known PUMA’s mechanism to induce cell death. These results indicate that the function of the individual components is preserved in the chimera. Our results point to a promising twofold strategy for drug delivery to; 1) avoid endocytosis by promoting liposome-membrane fusion, 2) provide additional protection by attachment to a stable, protease-resistant protein, which is a well-known method commonly used to prolong biologics half-life.

## INTRODUCTION

Proteins, peptides and their derivatives are used as therapeutic agents (biologics) in the treatment of a variety of medical conditions, including cancer, inflammatory diseases, genetic disorders and wound healing^1-3^. Some examples include hormones, enzymes, cytokines, antibodies and their fragments, and cell penetrating peptides^1-3^. The main advantages of biologics are their specificity, thus showing reduced levels of interference with non-targeted biological processes, and their affinity to natural chemical building blocks in the organism, thus reducing risk for toxicity. However, a major challenge for proteins, and particularly for therapeutic peptides, is their low efficacy due to short half-life resulting from proteolytic degradation caused by protease activity^1-3^. Therefore, biologics are rarely administered by ingestion or inhalation, but by intravenous or subcutaneous injection^4^. Despite which, many biologics still show very short half-life in circulation and often times are eliminated within minutes after administration, thus leading to treatments with frequent injections with the consequent patient discomfort and lack of compliance^1-4^.

This situation can be ameliorated by fusing the therapeutic peptide/protein to stable moieties of different chemical compositions, including oligosaccharides, polyethylene glycol (PEG), large unstructured polypeptides (XTEN) and plasma stable proteins such as albumin and IgG^5-7^. These strategies mainly consist in increasing the overall molecular size to protect the therapeutic from plasma protease degradation and decrease renal clearance, resulting in significantly increased half-life. However, another key elimination pathway of biologics is endocytosis, which involves the incorporation of the biologic to endosomes that eventually mature to late endosomes and lysosomes^8, 9^. Both organelles have internal acidic pH values and high concentrations of many different types of proteases, posing a great threat to biologics efficacy. To circumvent the endocytic pathway, a known methodology involves the fusion of the therapeutic to a protein that is recognized by endosomal receptors^6, 9^. The complex between the receptor and the fusion protein carrying the therapeutic is protected from degradation and stable under the endosomal acidic pH conditions. It is subsequently transported to the cell exterior, where it detaches from the receptor by the neutral pH of the extracellular milieu^6^.

Another successful strategy to protect biologics and increase their half-life is their encapsulation in liposomes, which can carry a variety of drugs with different chemical properties, including hydrophobic substances embedded in the lipid bilayer^10- 13.^Liposomes have been used to reduce the toxic effect of certain drugs by encapsulation during circulation, until the drug is released at specific target regions. Liposome lipid composition can be designed to modify bilayer characteristics, such as rigidity, overall charge, and to incorporate specific ligands. These modifications are typically aimed at targeting different types of cells. For example, the incorporation of cholesterol derivatives into electrostatically neutral liposomes helps them to resist circulation for several hours^14^. PEGylation of liposomes also helps decrease clearance by interfering with potential interactions between the liposome bilayer and proteins or other cell components in the reticuloendothelial system^15^.

Liposomes rarely fuse spontaneously with the plasma membrane bilayer to release their content to the cytosol, and typically enter the cell via endocytosis^13, 16^, thus leading to a pathway for potential degradation inside the endosomes and lysosomes. In this scenario, the use of an agent that can promote fusion of the lipid bilayer of the liposome with the cell membrane is particularly relevant^17^. Moreover, this agent could be a stable protease-resistant protein fused to a therapeutic peptide with the aim of increasing the half-life of the latter.

The protease-resistant, heat-stable, lysosomal protein saposin C (sapC), is a peripheral membrane protein with important roles in lipid degradation^18^ and lipid antigen presentation to CD1 (cluster of differentiation) proteins^19^. In the first case, sapC facilitates access of certain lipids (glucosylceramide) to the lysosomal hydrolase glucocerebrosidase. The molecular mechanism involves the binding of sapC both to the lipid bilayer and the enzyme, to assist the latter in lipid degradation^20, 21^. Mutations in either the enzyme or sapC result in the autosomal recessive disorder known as Gaucher disease caused by lipid accumulation in lysosomes^22^. In the second case, using likely an analogous mechanism, sapC associates to the lipid bilayer of endosomes and helps in the loading of lipid antigens to CD1 proteins^23^.

The three-dimensional (3D) structure of sapC in the absence of lipids was previously determined by NMR, revealing that sapC adopts the five-helix bundle motif with two pairs of helices connected by three disulfide bonds characteristic of the saposin fold (**Figure 1A**)^24^. This work also showed that the electrostatic surface of sapC is mainly negatively charged at neutral pH by the presence of abundant Glu and Asp amino acids in the protein surface (**Figure 1B**). SapC binds to lipids at acidic pH and detaches from the bilayer at neutral pH in a reversible process (**Figure 1C**)^24^. This behavior is explained by the neutralization at low pH of the negative charge of acidic residues at the protein surface, which otherwise preclude the protein from forming stable interactions with the hydrophobic tails of the lipids^24^. Further structural and biophysical studies revealed that the 3D structure of sapC in the presence of micelles conserves the secondary structure of the saposin fold, but the tertiary structure opens, exposing the hydrophobic core to detergent molecules (**Figure 1D**)^25^.

**Figure 1.**
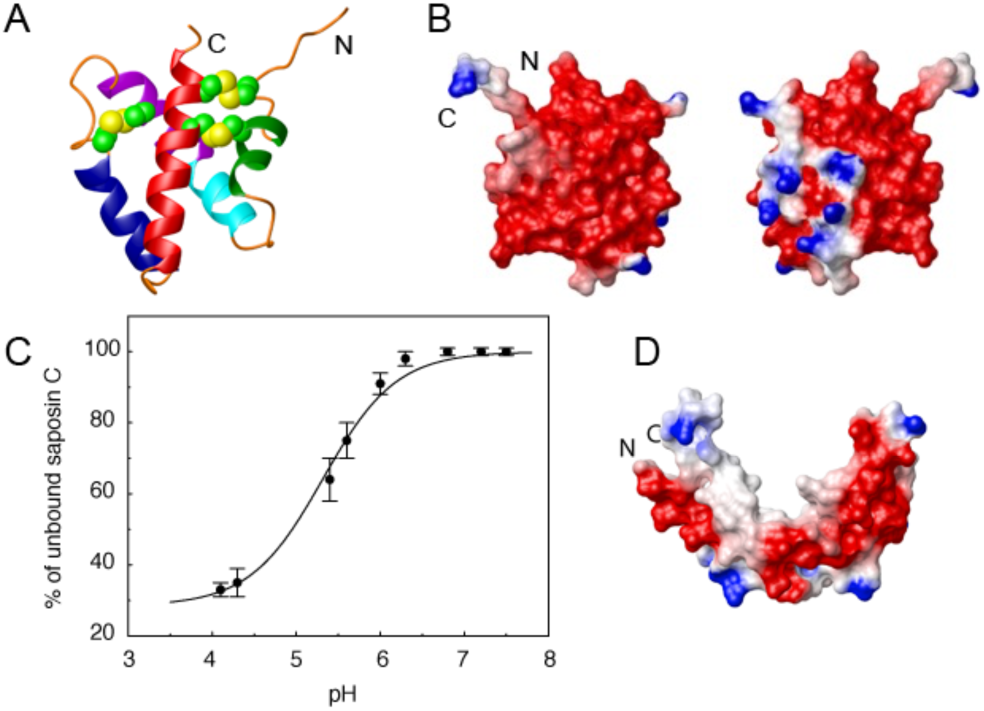
SapC binding to liposomes is a pH-dependent reversible process. A. Ribbon diagram of the 3D NMR structure of sapC^24^. B. Electrostatic surface charge distribution of sapC. C. SapC reversible binding to liposomes with pH. D. 3D structure of sapC bound to micelles^25^.

Importantly, sapC has been reported to produce liposome fusion at acidic pH^26, 27^, although the specific molecular mechanism is not understood.

All in all, sapC shows special characteristics, including high stability and protease resistance, liposome fusogenic activity and pH-dependent liposome binding. Moreover, sapC can be over-expressed by recombinant methods and purified by standard chromatographic techniques. These attributes point to sapC as an attractive candidate for biotechnological applications. In fact, sapC bound to liposomes formed by the phospholipid 1,2-dioleoyl-sn-glycero-3-phospho-L-serine (DOPS) has been shown to target cancer cells and promote apoptosis when injected intravenously in mice^28, 29^.

The method to prepare the sapC-DOPS complex consists in mixing the protein solution with lipids at pH ∼ 5^29^. This solution is then diluted with a buffer to reach physiological pH. Once liposomes are formed at pH ∼ 5, sapC will be bound to the inner and outer leaflets of the liposome bilayer. However, once sapC-DOPS is in contact with a fluid/solution at physiological pH, sapC bound to the outer leaflet will be released. This result has been confirmed by different laboratories^23^, including the authors of the sapC-DOPS particle (reference 29 Suppl. Material), and matches our previous structural studies on the pH dependence of sapC binding to liposomes and micelles^24^. The mechanism for this specific targeting of cancer cells by sapC-DOPS has been proposed to involve the preferred binding of sapC to the high levels of phosphatidyl serine (PS) in the outer leaflet of cancer cell membranes^28^. However, intravenous injection of sapC-DOPS will result in the detachment of sapC from the liposome outer leaflet due to the slightly basic blood’s pH value (pH ∼ 7.4). Therefore, further studies are necessary to investigate the mechanism of recognition of sapC-DOPS and cancer cells.

We show here that sapC in combination with liposomes can be merged to therapeutic peptides with a twofold purpose; i.e. to confer stability to the therapeutic in the presence of extracellular proteases, and to promote membrane-liposome fusion that can potentially avoid the endocytic pathway and therefore decrease the chance of degradation inside the cell. As proof of concept, we have designed and characterized at the functional level a protein chimera carrying sapC at the N-terminus and the active domain (BH3 domain) of the Bcl-2 proapoptotic protein PUMA, a ∼ 25 amino acid-long peptide covalently attached to it by recombinant methods (sapC-PUMA).

PUMA is an intrinsically disordered protein of the BH3-only subfamily that promotes apoptosis by antagonizing prosurvival Bcl-2 members^30^. The known molecular mechanism that PUMA follows to induce apoptosis consists in direct binding to antiapoptotic Bcl-xL^31^. It has been shown that the ∼ 25 amino acid-region spanning PUMA’s BH3 domain (PUMA^BH3^) suffices for promoting cell death. BH3-derived peptides and peptidomimetics are intensively investigated as anticancer drugs^32, 33^. Some BH3 domains are specific for certain prosurvival proteins; however, proapoptotic PUMA^BH3^ is promiscuous and po-tently binds with high affinity multiple prosurvival Bcl-2 proteins^34^. Because of these special characteristics, PUMA^BH3^ is an ideal candidate for peptide-based anticancer drug.

We show by NMR that the structure of sapC is unperturbed by the presence of the covalently attached PUMA^BH3^ peptide within the chimera (sapC-PUMA), and it is still capable of binding liposomes under mildly acidic conditions (pH 6.0). Analogously, our NMR data show that PUMA^BH3^ in the chimera is capable of binding Bcl-xL. The binding affinity of sapC-PUMA to liposomes is low at pH 7, which is not optimal for the liposomes to carry the therapeutic in the outer leaflet. With the purpose of enhancing sapC-PUMA binding to liposomes at neutral pH, we have designed and studied a mutant of the chimera by substituting two acidic residues in the sapC domain by positively charged amino acids. We show that liposome binding increases at pH 7 for the mutant chimera compared to the wildtype, thus pointing to a suitable strategy for additional modifications to enhance binding at neutral pH.

## RESULTS

### Design of protein chimeras and protein solubility tests

The sequence of the sapC-PUMA chimera is shown in **Figure 2**. Human sapC is an 80 amino acid-long protein to which a short linker of three Gly amino acids is attached at the C-terminus to connect 25 additional amino acids encompassing PUMA^BH3^. The few extra amino acids at the C-terminus of the chimera include the His-tag for purification by affinity liquid chromatography, following a thrombin cleavage site to subsequently remove the His-tag. We opted to use a short linker connecting the two components, because a previous chimera design with a ∼ 20 amino acid-long linker resulted in the absence of binding of sapC to liposomes (data not shown). We speculate whether lack of binding is a result of steric hindrance impeding sapC from open its structure for liposome interaction.

**Figure 2.**
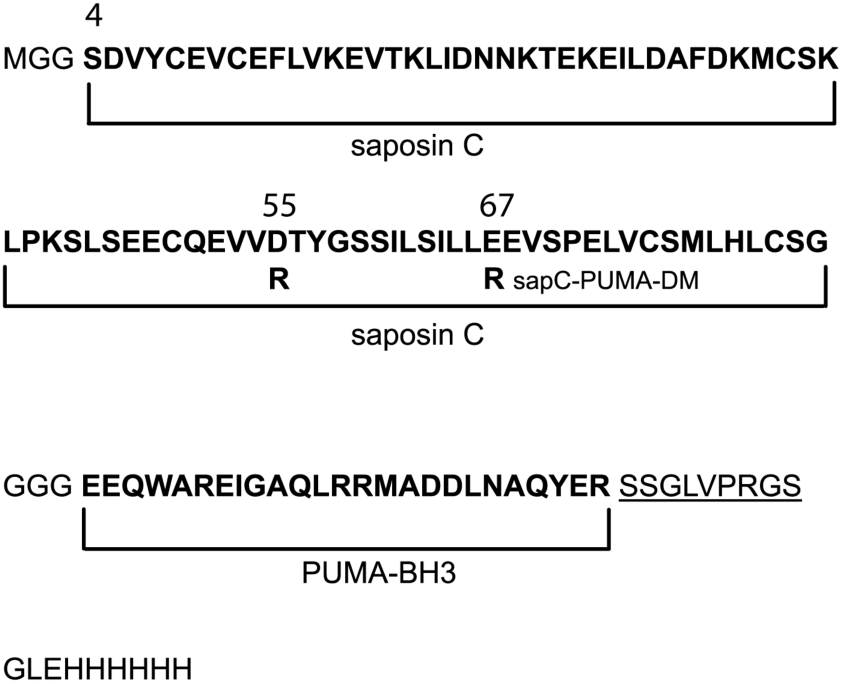
Amino acid sequence (one-letter code) of engineered protein chimera sapC-PUMA and double-mutant; sapC-PUMA-DM (D55R, E67R, mutations indicated in the figure). The regions corresponding to sapC and PUMA^BH3^ are denoted in bold. The thrombin cleavage site is underlined followed by the His-tag. The natural N-terminus amino acid of sapC is labeled as “S4”. Other numbers indicate the position of the mutations.

Previous studies on a conservative mutant of sapC (Glu 6 Gln, Glu 9 Gln; original numbering) designed to remove two negative charges from the protein surface, show that the mutant binds to liposomes to a greater extent under identical conditions as compared to wildtype sapC^24^. Based on this information, and with the aim of increasing liposome binding of sapC-PUMA chimera at neutral pH, we designed a non-conservative double-mutant (sapC-PUMA-DM) that replaces two negatives charges by mutating amino acids Asp 55 and Glu 67 to Arg (**Figure 2**). These mutations were chosen because of their location far from the contact region of sapC to the lipid bilayer based on the structure shown in Figure 1D^25^.

Because the binding of sapC to liposomes is pH-dependent, we studied the solubility of the different chimeras in a broad pH range to identify the best compromise between liposome biding and protein solubility. This information will be necessary to design different applications of sapC chimeras/liposome assemblies. Our results indicate that the isoelectric point of each chimera increases as the number of mutations to positively charged amino acids increases, as expected, and we observe that the overall solubility of the mutant chimera also increases (**Figure 3**).

**Figure 3.**
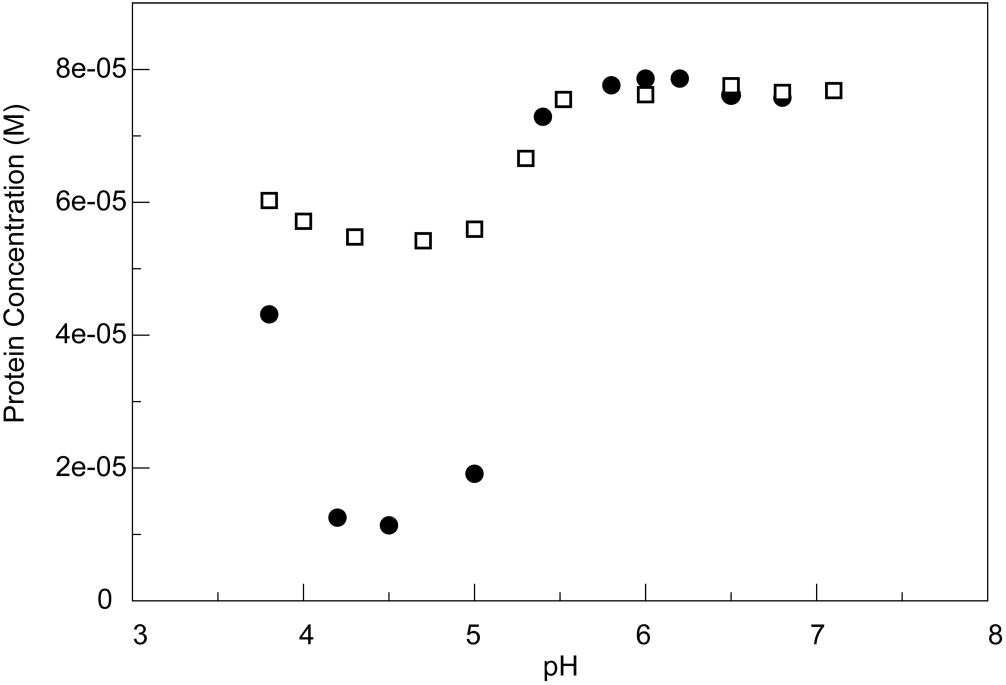
The solubility of the chimera increases by engineering the electrostatic surface of sapC. SapC-PUMA (circles) and sapC-PUMA-DM (squares) solubility with pH measured by absorbance spectroscopy at 280 nm.

### Function of the sapC domain in the chimera construct

To test whether the presence of PUMA in the sapC-PUMA chimera affects sapC at the structural level, we acquired [^1^H-^15^N]-2D NMR spectra of the chimera to compare with wildtype sapC. This type of experiment is known in protein NMR as the “protein fingerprint” because it will be significantly perturbed upon small conformational or structural changes in the protein. The NMR data indicate a high degree of overlap between the ^1^H-^15^N amide signals of wildtype sapC and sapC within the chimera, which allowed almost the full assignment of the amide ^1^H and ^15^N chemical shifts of the sapC component in the chimera (**Figure 4**). As expected, the [^1^H-^15^N]-2D spectrum of sapC-PUMA shows additional signals that correspond to the linker and PUMA^BH3^ (**Figure 4**). These results indicate that the structure of sapC within the engineered protein is unperturbed.

**Figure 4.**
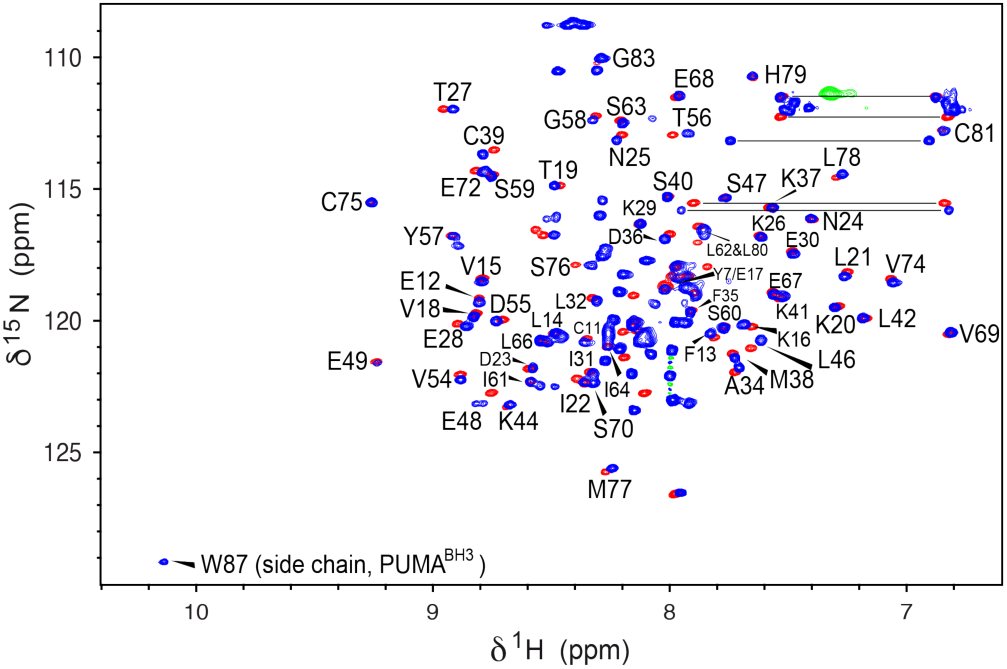
SapC overall structure is retained in sapC-PUMA. [^1^H-^15^N]-HSQC NMR spectra of sapC (red) and [^1^H-^15^N]-sofast_HMQC^35^ of sapC-PUMA (blue). Labels indicate amino acids in sapC-PUMA that do not show significant changes in chemical shifts. The NH signal of the Trp indole ring in PUMA is labeled. Several side chain NH2 signals of Asn and Gln residues are connected with a horizontal line.

The combined amide ^15^N and ^1^H chemical shift deviations between sapC and sapC-PUMA versus the amino acid sequence are shown in **Figure 5**. Some amino acids show larger deviations probably due to slight changes in the solution pH (**Figure 5**). For example, Glu 48, is one of the signals with the largest chemical shift changes. Amino acids Pro 43 and Pro 71 are not observed because they lack the NH amide pair that gives rise to signal in [^1^H-^15^N]-2D spectra. Several amino acids such as Gln 51, Glu 52, Ile 64 and Leu 65, could not be unambiguously as-signed and are not shonw. Altogether, the average chemical shift change observed is 0.03 ± 0.02 ppm.

**Figure 5.**
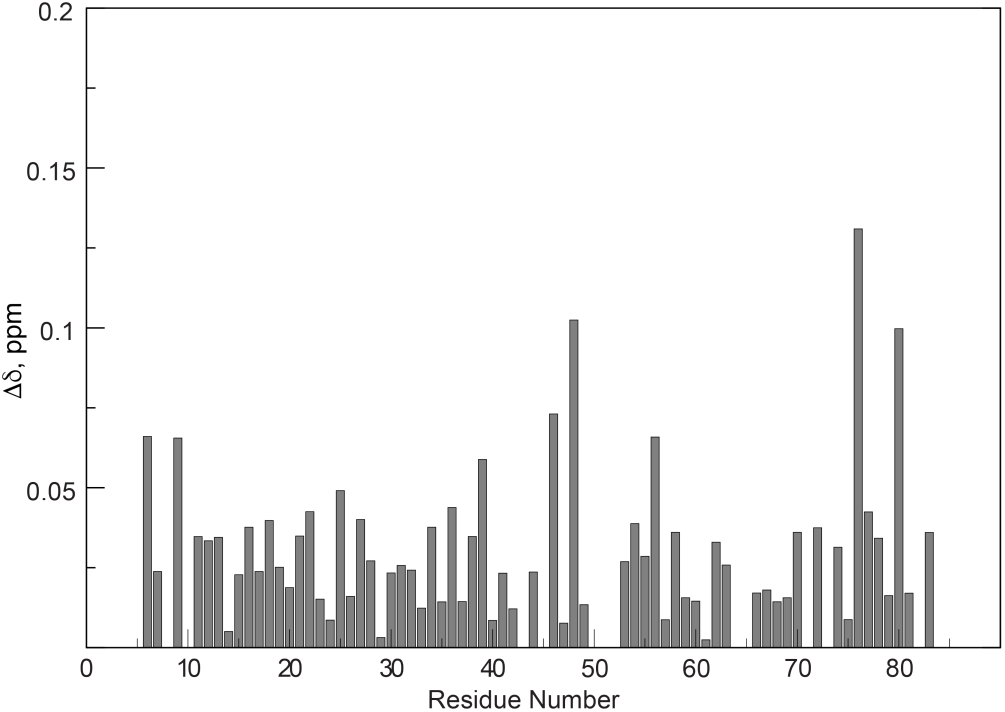
Small chemical shift differences between free sapC and the sapC domain of sapC-PUMA. The combined amide ^1^H and ^15^N chemical shifts were calculated with the equation:

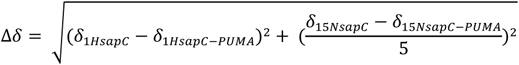

Previously, we showed that sapC is capable of binding liposomes composed of phosphatidyl serine (PS) and phosphatidyl choline (PC)^24^. The binding at 1:1 protein/lipid molar ratio is negligible at neutral pH but increases at more acidic pH following a sigmoidal behavior, which reaches a plateau with binding close to 70 % at pH 4.2 (**Figure 1C**)^24^.

Based on the absence of structural changes in the sapC domain of the chimera, it is reasonable to expect that the function of sapC is also unmodified. To prove this hypothesis, we performed liposome binding assays with sapC-PUMA using NMR. Because of the low solubility of the chimera at pH < 5.8, we carried out the titrations at pH 6.0 with increasing values of the lipid concentration. SapC is known to show preferred binding for negatively charged lipids such as PS^26^. Thus, for the titration experiments at a pH 6 (conditions for low liposome binding), we used PS liposomes. One-dimensional projections of [^1^H-^15^N]-sofast-HMQC experiments^35^ on the chimera-liposome mixtures show a significant decrease in the NMR signal intensity at increasing concentrations of lipid (**Figure 6**). This result is analogous to that observed for the binding of wildtype sapC to PS/PC liposomes^24^. The large size of the liposomes (∼ 100 nm) results in an overall particle tumbling time that is very slow compared to that of the free protein (4.6 ns)^24^, which significantly impacts NMR signal to noise ratio due to magnetic relaxation processes. In fact, very large particles are “invisible” in NMR. Thus, the overall NMR signal intensity decreases when the protein binds to liposomes because the effective population of protein free in solution decreases (**Figure 6**).

**Figure 6.**
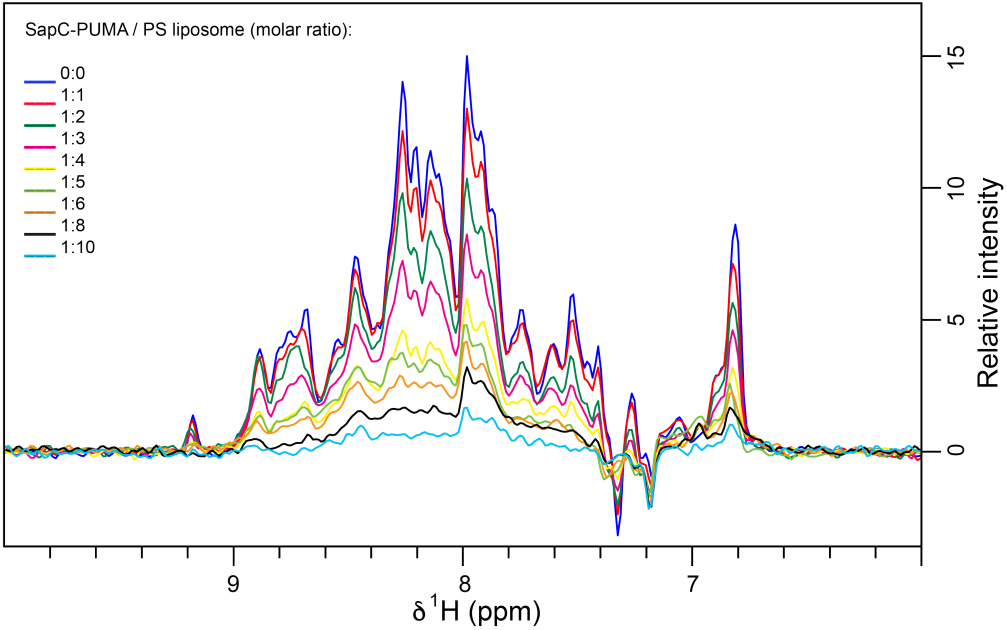
SapC-PUMA binds to liposomes at pH 6. NMR signal intensity gradually decreases as the concentration of liposomes increases due to proteinlipo-some binding.

### The binding of sapC-PUMA to liposomes can be tuned by modifying the electrostatic surface of sapC

The decrease in NMR signal intensity in the liposome titration follows a binding isotherm that allows to estimate the dissociation constant (K_D_) for the binding of sapC-PUMA to PS liposomes (**Figure 7**). SapC-PUMA binds liposomes with an apparent K_D_ of 757 μM at pH 6.

**Figure 7.**
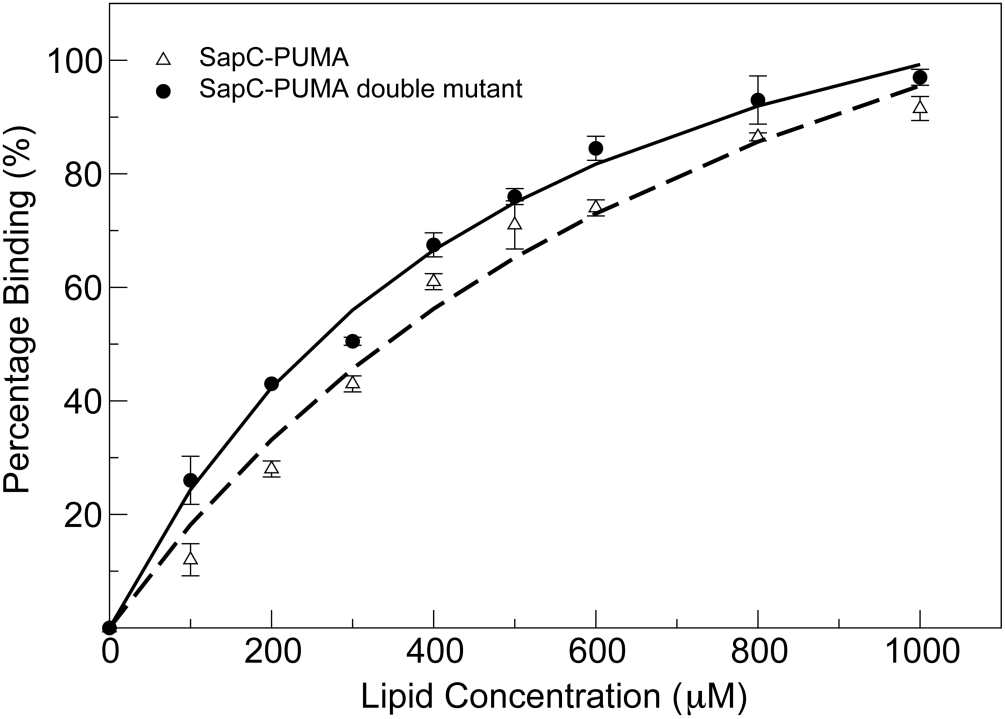
SapC-PUMA affinity for liposomes is increased by modifying the electrostatic surface of sapC. Percentage binding of sapC-PUMA and sapC-PUMA-DM are shown with triangles and circles, respectively. Measurements were done in duplicate and the corresponding error bars are indicated.

The mutant chimera (sapC-PUMA-DM) in which the electrostatic surface of the sapC domain is more positively charged due to mutations of Asp 55 and Glu 67 to Arg, is expected to bind with increased affinity to liposomes. We firstly checked using [^1^H-^15^N]-2D NMR that the saposin fold was not perturbed by the presence of the mutations in the sapC domain. **Figure SM1** shows an overlay of the spectra of sapC-PUMA and sapC-PUMA-DM, indicating minimal perturbations of the chemical shifts and a properly folded protein. We then followed the same procedure for the NMR titration of sapC-PUMA-DM with liposomes. Our data indicate that liposome binding is enhanced in the presence of additional positive charges in the surface of sapC, as the apparent K_D_ for the binding of sapC-PUMA-DM to PS liposomes is 399 μM at pH 6. Thus, the affinity of sapC-PUMA-DM for liposomes has almost doubled relative to wildtype sapC-PUMA (**Figure 7**).

We also tested the binding of the different chimeras in mixtures at 1:10 protein/lipid molar ratio at pH 7 for potential therapeutic applications of the sapC-PUMA/liposome assemblies at physiological pH. As expected, binding is low for the wildtype chimera, showing a percentage of protein bound to liposomes from NMR data of 8.5%, whereas it increases to 28% for sapC-PUMA-DM (**Figure SM2**). These data indicate that amino acids that are positively charged in the surface of sapC facilitate liposome binding at physiological pH.

Our results have significant implications for sapC-DOPS particles currently being used to target cancer cells^29^. The intravenous injection of these assemblies will result in the removal of wildtype sapC from the outer leaflet of the liposome, thus compromising the recognition of specific cell populations. However, sapC mutants with increased positive charges in the protein surface, like sapC-PUMA-DM, will remain attached to the outer leaflet of the liposome at neutral pH, potentially enhancing cell targeting. Altogether, our results indicate that liposome binding can be tuned by modifying the electrostatic surface of sapC, which could result in different therapeutic applications of the protein/ liposome assembly.

### Function of the PUMA domain in the chimera constructs

PUMA is a BH3-only protein (containing the Bcl-2 homology domain #3) of the Bcl-2 family that promotes apoptosis or programmed cell death^34^. Pro- and anti-apoptotic members of the Bcl-2 family participate in protein-protein interactions as a mechanism to regulate cell fate^36^. It is well known that the BH3 region is the active biding component of the proapoptotic proteins. Thus, peptides comprising the BH3 domain are capable of binding and eliciting cell death^30^. In addition, it has been extensively reported that certain types of cancer cells are resistant to anticancer treatments due to mechanisms developed to impair apoptosis^37^. Therefore, BH3-derived peptides from proapoptotic members of the Bcl-2 family are intensively investigated as anticancer drugs^38^. The BH3 domain of PUMA is capable of binding the prosurvival protein Bcl-xL as a mechanism of antagonizing its function and thus promoting cell death^34^. However, PUMA is unique in that it is the only protein capable of destabilizing the interaction between Bcl-xL and the transcription factor p53 to induce cell death^39^. Once free, p53 can activate proapoptotic Bax and Bak, which trigger apoptosis by mitochondrial outer membrane permeabilization^39^. The complex between PUMA^BH3^ and Bcl-xL has been determined by NMR and the K_D_ for PUMA^BH3^ and Bcl-xL binding is ∼ 3 nM^31^.

To test the binding of the PUMA^BH3^ domain in the sapC-PUMA chimera, we have performed NMR titration experiments on ^15^N-labeled sapC-PUMA with unlabeled Bcl-xL at three different molar ratios (sapC-PUMA/Bcl-xL: 1:0.5, 1:1, 1:2). The binding of sapC-PUMA to Bcl-xL is apparent from the overall decrease in signal intensity and disappearance of numerous signals in the spectrum of the complexed chimera compared to free sapC-PUMA. The decrease in signal intensity as the concentration of Bcl-xL increases is plotted for some amino acids of the sapC domain in **Figure 8A**. This result can be explained by the increase in molecular weight (slower tumbling rate) once sapC-PUMA (∼14.4 kDa) binds Bcl-xL (∼24.5 kDa). In addition, large chemical shift changes are observed for signals that belong to the PUMA^BH3^ domain (**Figure 8C&D**), whereas sapC signals are minimally perturbed (**Figure 8B**), indicating that the former is the domain binding to Bcl-xL.

**Figure 8.**
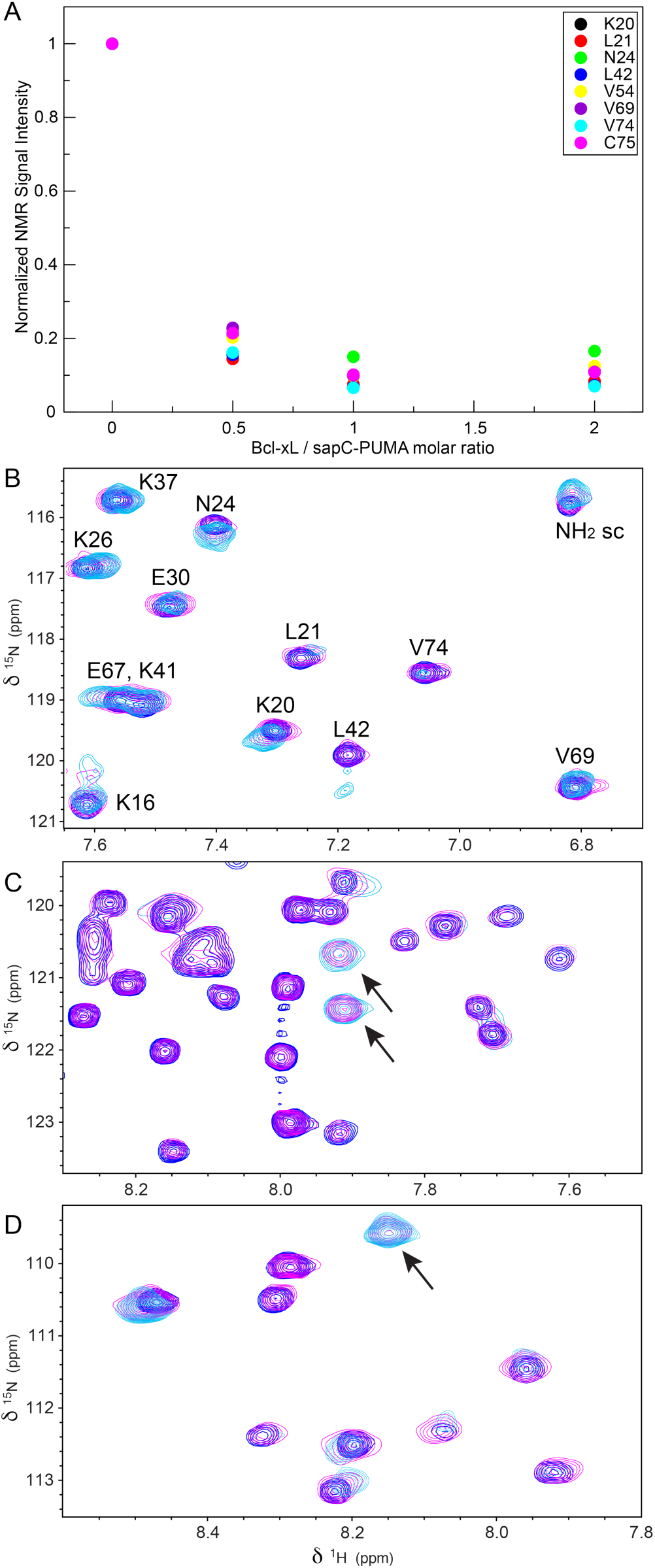
SapC-PUMA binds Bcl-xL via PUMA^BH3^ domain. A. NMR signal intensity decrease for selected amino acids (indicated in the inset) in sapC-PUMA as Bcl-xL/sapC-PUMA molar ratio increases. B, C, D. Selected regions of overlaid [^1^H, ^15^N]-2D NMR spectra of ^15^N-labeled sapC-PUMA at Bcl-xL/sapC-PUMA molar ratios; 0:1 (blue), 0.5:1 (magenta) and 1:1 (cyan). SapC signals labeled with the corresponding amino acids are minimally perturbed in B. New signals for the PUMA domain (indicated with arrows in C and D) are observed at Bcl-xL/ sapC-PUMA molar ratios 0.5:1 and 1:1.

The spectrum at 0.5:1 molar ratio of Bcl-xL/sapC-PUMA displays two sets of signals for the PUMA^BH3^ domain corresponding to populations of the unbound and bound conformations (signals of the bound conformation indicated by arrows in **Figure 8C&D, magenta**). These signals are not observed in the spectrum at 0:1 molar ratio (no bound population) and are still observed in the spectrum at 1:1 molar ratio (100% bound population). When the Bcl-xL/sapC-PUMA molar ratio is increased to 1:1 or 2:1, only one set of signals remains, which corresponding to 100% population of bound PUMA^BH3^ (**Figure 8 C&D**). Overall, these results indicate that the sapC-PUMA/Bcl-xL complex is 1:1 and that the exchange rate between the bound and unbound conformations of sapC-PUMA is slow on the NMR chemical shift time scale as two sets of signals are observed.

Because the binding of sapC-PUMA to Bcl-xL is in the slow exchange regime, it is not possible to calculate a K_D_ value (K_D_ = k_off_ / k_on_) from the changes in chemical shifts or signal intensity upon binding. The former do not represent population averages of the free and bound forms as in the case of fast exchange, and the latter are affected by the very different relaxation times of the free and bound species in this case. However, it is possible to estimate an upper limit of the K_D_ value based on the differences in chemical shifts of the two sets observed. The difference in amide ^1^H chemical shift of PUMA signals in the free and bound forms (∼ 0.082 ppm) (**Figure 8C**) results in a value for Δω of ∼ 310 rad s^-1^ at the spectrometer frequency of 601.13 MHz used for these studies. Assuming that binding is diffusion-controlled; k_on_ ∼ 10^9^ M^-1^ s^-1^ and k_off_ ∼ 10^9^. K_D_. The slow exchange regime observed indicates that k_off_ << 310 s^-1^ and therefore K_D_ << 310 nM, as expected based on previously reported values for the binding of PUMA^BH3^ and Bcl-xL.

We have also checked by NMR that the mutant chimera, sapC-PUMA-DM binds Bcl-xL with similar upper value of the dissociation constant, as we observe two sets of signals indicative of slow exchange regime in the spectrum of sapC-PUMA-DM upon titration with Bcl-xL (**Figure SM3**).

### Liposome fusion orchestrated by sapC

Experiments reported previously to prove the fusogenic capabilities of sapC were done using TEM^26^, SEC^26^ and dynamic light scattering^27^ under certain conditions that do not undoubtedly point to vesicle fusion. For example, comparative TEM micrographs of negatively stained liposome samples in the absence and presence of sapC have been used to interpret vesicle size increase. Negatively stained TEM grids need to be completely dry before image acquisition; a process that can distort liposome size and shape. To illustrate this fact, we have obtained TEM micrographs of negatively stained liposome samples prepared using different procedures, showing the variability of size and shape distortion (**Figure SM4**). Size-exclusion chromatography (SEC) is commonly used to determine the size distribution in liposome preparations and to separate empty from loaded liposomes^40^. However, it is well known that liposomes interact with the SEC matrices in a dynamic and reversible manner, which results in misleading retention times^41^. To avoid this effect, it is necessary to initially coat the SEC matrix with lipids, which was not done in reported experiments^26^. We attempted to monitor liposome size using SEC and found that consecutive injections of the same liposome preparation resulted in chromatograms with peaks at different elution volume (data not shown), thus rendering unreproducible results.

Dynamic light scattering is an ideal technique to investigate liposome size. This technique has been used previously to test liposome fusion mediated by sapC using a protein construct carrying the His-tag^27^. This work reports an increase in liposome size from ∼ 200 nm to 3 μm. However, we found using NMR that sapC-PUMA with His-tag shows high tendency to aggregate, as observed by severe line broadening in the ^1^H NMR spectrum when the pH is decreased from 6.8 to 4.2 (conditions necessary for abundant liposome fusion) (**Figure SM5**). In contrast, previously reported NMR studies on sapC without His-tag indicate that the protein is monomeric at acidic pH^24^. Thus, we hypothesized that sapC with His-tag could show significant aggregation at acidic pH, which could interfere with the dynamic light scattering results reporting on liposome fusion.

We have used dynamic light scattering to study liposome fusion orchestrated by sapC (without His-tag) and its time dependence. We observed that in approximately 5 minutes after addition of sapC to liposome preparations, liposomes increase in diameter from ∼ 100 nm to ∼ 400 nm and reach approximately 900 nm after 200 minutes (**Figure 9**). Our results indicate that the increase in size is significantly smaller than previously reported^27^.

**Figure 9.**
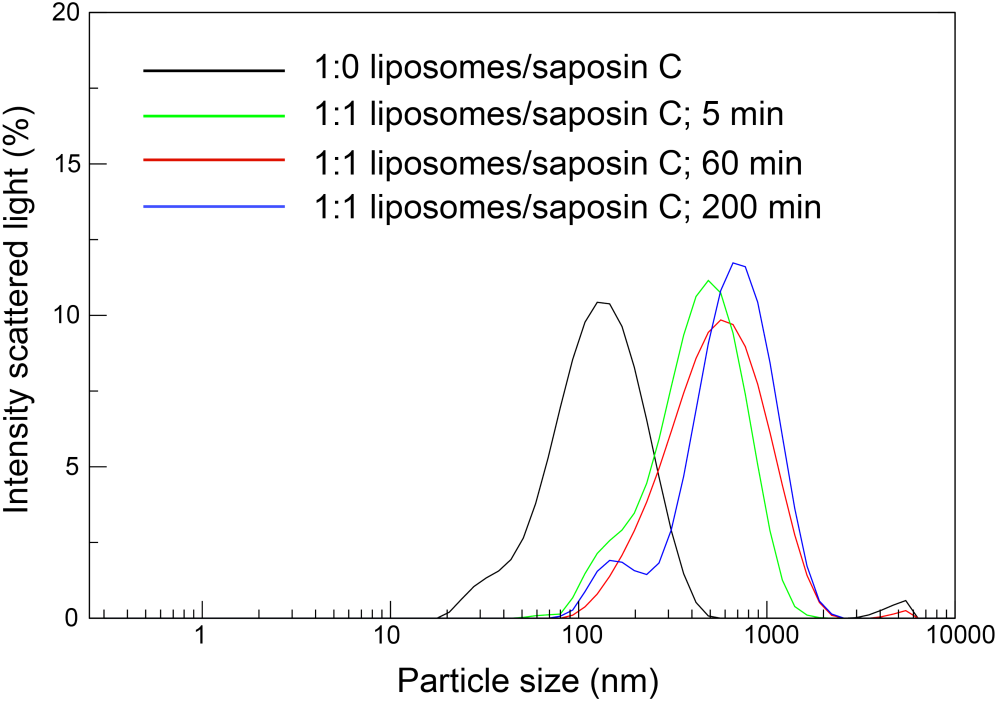
Liposome fusion mediated by sapC. The analysis of the intensity of the scattered light reveals an increase in the diameter of liposomes in a time-dependent manner after the addition of sapC.

Our dynamic light scattering data clearly show that sapC can fuse liposomes, leading to an approximately 10-fold increase in liposome diameter. This result, combined with the proven capability of sapC-PUMA to bind liposomes, suggests that sapC can be used in therapeutic protein chimeras to leverage the fusogenic capabilities of sapC to merge liposomes to plasma membranes. This effect could have a significant impact in the stability of biologics by protecting them from endocytic pathways.

## CONCLUSION

We have designed the chimera protein sapC-PUMA as proof of concept of a twofold strategy for biologics protection from degradation. The sapC domain has two functions: 1) to protect the therapeutic component of the chimera, PUMA is this case, by increasing the overall size of the biologic with a protease-resistant and stable protein; 2) to allow the incorporation of the biologic into liposome nanoparticles that will promote the fusion between the liposome and the plasma membrane, thus avoiding the incorporation of the biologics via endocytosis. In the event that the sapC chimera enters the cell by endocytic pathways, the specific characteristics of sapC as a lysosomal, acidic pH- and protease-resistant protein, would confer additional protection to the biologic.

We show by NMR that both domains (sapC and PUMA^BH3^) in the chimera are still functional at the molecular level. SapC-PUMA retains the saposin fold and sapC’s functionality by exhibiting reversible pH-dependent binding to liposomes. In addition, the PUMA^BH3^ component of the chimera binds tightly to Bcl-xL, its natural antagonist, in the presence of the sapC domain. These results indicate that both sapC and PUMA^BH3^ have minimal interference in each other’s function, thus fulfilling the most basic requirement of the chimera design.

Previous work reported that maximum binding of sapC to liposomes occurs upon acidification^24^. Here we show that binding can be observed and quantified at mildly acidic conditions (pH 6) by significantly increasing the lipid to protein molar ratio. Importantly, we demonstrate that it is possible to modify the affinity of the protein to liposomes by changing the overall charge of sapC’s electrostatic surface in the chimera. These modifications serve a twofold purpose; 1) increase the overall solubility of the chimera while retaining the saposin fold and 2) allow to tune the pH dependence of liposome binding. With these two mutations replacing acidic for basic amino acids, the chimera almost doubles its binding affinity to liposomes.

In addition, the significant increase in liposome binding of the chimeric mutant at pH 7 points to the possibility of keeping sapC chimeras bound to the liposome outer leaflet at physiological pH for different biomedical purposes. These results are particularly relevant to the applications of sapC-DOPS, which is formulated using wildtype sapC with negligible liposome binding at physiological pH^23, 24, 29^. Our results on the pH-tuning of liposome binding by sapC mutation open the door to potentially target tissues at different pH values depending on the need, as the formulation of sapC chimera-liposomes can be altered for optimal interaction. This possibility is analogous to current research efforts directed to target tumors and control drug release at different pH values by proper liposome functionalization^42^.

The sapC-PUMA chimera with apoptotic function holds promise as an anticancer drug with the ability to be delivered intravenously while bound to liposomes, thus protecting it from protease degradation. Altogether, our results indicate that sapC is a good candidate for engineering chimeric biologics by adding peptides or small proteins as potential therapeutics. However, the size of the therapeutic and the length of the linker connecting both components needs to be carefully designed, as our prior attempts with longer linker and large peptide inhibited sapC binding to liposomes.

Future work is necessary to test the effect of other therapeutic peptides/proteins in sapC chimeras at the molecular and functional levels, and to further tune the liposome binding affinity with additional mutations to target specific tissues and promote liposome fusion under different pH conditions.

## EXPERIMENTAL SECTION

### Protein purification

The DNA sequences of sapC-PUMA and sapC-PUMA-DM were inserted in pET-30b vectors containing a thrombin cleavage site at PUMA’s C-terminus. Purchased plasmids (Gene Universal) were transformed into *Escherichia coli* BL21(DE3) cells and grown in ^15^N enriched minimal media for NMR studies and in Luria Broth for all other experiments. Protein expression was induced using 1 mM isopropyl β-D-thiogalactopyranoside (IPTG) at 37 °C for 4 hours. Cells were harvested by centrifugation at 8K rpm for 30 minutes and resuspended in 20 mM Tris (pH 8.0), 20 mM imidazole, 500 mM NaCl, and phenylmethylsulfonyl fluoride (PMSF) as protease inhibitor. Cells were broken by sonication at 20 kHz for 48 minutes on ice, with sonication at intervals of 15 sec with resting periods of 45 sec and centrifuged at 35 k rpm for 45 min. Cell lysate was filtered through a 0.45 µm pore filter. Protein constructs contained a C-terminal six-His tag were purified using Ni^2+^ affinity chromatography. Proteins were eluted using an HPLC gradient of 500 mM imidazole. Afterwards, the proteins were dialyzed using 20 mM Tris (pH 8.0), 150 mM NaCl, 2.5 mM CaCl_2_, and subjected to thrombin cleavage overnight. For 1D-^1^H NMR experiments (*vide infra*), some protein samples were not cleaved to retain the His-tag. In all cases, the proteins were further purified by reverse phase chromatography using a C4 column (Higgings An-alytical) in water and acetonitrile mixtures with 0.1% TFA. Proteins were lyophilized after reverse phase. The solid protein was then resuspended in HPLC-grade water for different experiments.

Bcl-xL-ΔTM (without the transmembrane domain) was cloned into the pET21a vector with a N-terminal six-His tag and was purified using the same methods described for sapC-PUMA constructs, with the addition of 500 µM tris (2-carboxyethyl) phosphine (TCEP) in the resuspension buffer and elution buffer for Ni^2+^ affinity chromatography. Bcl-xL-ΔTM was not subjected to dialysis or thrombin cleavage before reverse phase chromatography.

### Liposome preparation

Brain L-α-phosphatidylserine lipids were purchased in chloroform from Avanti Polar Lipids. Lipids were dried overnight in vacuum and resuspended in water to a lipid concentration of 2 mM. Lipids were allowed to hydrate for 1 hour, vortexed for 10 minutes, and sonicated by bath sonication for 20 minutes. Ice was added to the bath sonicator to control the temperature during sonication. Liposome size was determined using dynamic light scattering.

For TEM preparations liposomes at 1 mM concentration were dried in vacuum, dissolved in 20 mM acetate buffer, vortexed and bath sonicated. After sonication the liposomes were subjected to ten cycles of extrusion through a polycarbonate filter with a pore of 100 nm.

Other liposome preparations for Size Exclusion Chromatography and dynamic light scattering experiments include the hydrating and vortexing steps mentioned above with the following modifications: 1) use of tip sonication at 4 kHz for 6 min (30 sec on, 30 sec resting) on ice; 2) use of tip sonication followed by extrusion through either 100 nm or 200 nm pore filter; 3) same as 2) with the addition of 10 freeze-thaw cycles following the first extrusion set, and a second 20-time extrusion after the freeze-thaw cycle. Freeze-thaw cycles include 3 minutes in dry ice bath with ethanol, followed by 3 minutes in 50 °C water bath. Data for these preparations are not shown.

We found that liposomes that were tip sonicated or bath sonicated were consistent in size with a diameter of approximately ∼ 100 nm. Liposomes that underwent freeze-thaw cycles, whether they were extruded through 100 nm or 200 nm pore filter, resulted in an average diameter close to 190 nm based on dynamic light scattering data.

### Solubility studies with pH

Solubility studies were conducted using an Agilent Cary 60 UV-Vis spectrophotometer at 280 nm. Prior to each measurement the protein was centrifuged to pellet any particles and the supernatant was filtered through a 0.22 μm filter. The cuvette was washed between measurements and a blank measurement was run to ensure no residual protein material remained attached to the walls. SapC-PUMA and sapC-PUMA-DM solutions at 100 μM were initially brought at neutral pH, and the pH was subsequently decreased by incremental addition of dilute solutions of HCl. Data were analyzed using QtGrace.

### NMR spectroscopy

NMR experiments were acquired on a Bruker Avance III 600 MHz spectrometer equipped with a cryoprobe. All samples were prepared at 10% D_2_O, 90% HPLC-grade H_2_O. 2D-NMR data were collected at 298 K using [^1^H-^15^N]-SOFAST-HMQC experiments^35^. Amide ^1^H-^15^N chemical shift assignments for sapC in the sapC-PUMA chimeras were obtained using previous assignment of sapC^24^. The spectra of sapC-PUMA had to be shifted in the ^1^H and ^15^N dimensions relative to sapC due to overall displacement of signals when overlaying the spectra. This could be due to differences in the temperature calibration of the different probes used and to differences in the calibration of the base frequencies of ^1^H and ^15^N of the different spectrometers. PUMA^BH3^ was not assigned, except for the NH moiety of the Trp side chain. Data were processed with TOPSIN and NMRPipe software^43^ and analyzed with SPARKY^44^.

### NMR sample preparation for sapC-PUMA and liposome binding experiments

Initial liposome binding studies were done using 1D-^1^H NMR in unlabeled sapC-PUMA constructs with and without the His-tag prepared to a final protein concentration of 100 µM at pH 6.8 and 100 µM lipid concentration of liposomes. Samples with and without liposomes were subjected to pH adjustments using dilute solutions of HCl and NaOH in the pH range from 6.8 to 4.2. All samples were inserted into clean NMR tubes to avoid minor pH changes. 1D-^1^H NMR was used to confirm liposome binding at acidic pH for constructs without His-tag. Constructs with His-tag showed extensive line broadening at acidic pH in the absence of liposomes indicating significant protein aggregation (**Figure SM5**). Thus, all experiments for liposome titration were done on samples without the His-tag.

Liposome titrations were done using [^1^H-^15^N]-sofast-HMQC^35^ experiments on ^15^N labeled sapC-PUMA and sapC-PUMA-DM samples. Stock solutions were prepared in HPLC-grade water at pH 6.0 and 500 µM protein concentration, confirmed by absorbance at 280 nm. The final concentration of sapC-PUMA used for NMR experiments was 100 μM at pH 6.0. ^15^N SapC-PUMA constructs were mixed with increasing lipid concentration of PS liposomes at pH 6.0. The following sapC-PUMA:lipid molar ratios were tested in sequential order: 1:0, 1:1, 1:2, 1:3, 1:4, 1:5, 1:6, 1:8 and 1:10. The first 1D projections of the 2D NMR experiments were used to monitor signal intensity decrease during the liposome titration. All 1D projections were baseline-corrected from 11 ppm to 4.7 ppm. The signal intensity was obtained by integration in the range from 9.3 ppm to 6.6 ppm using TOPSPIN software. The binding isotherms were fitted using Grace to the equation:

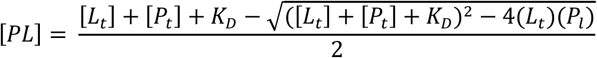

Where [PL] is the concentration of protein-lipid complex, [L_t_] is the total concentration of lipid, [P_t_] is the total concentration of protein and K_D_ is the dissociation constant of the protein-lipid complex.

### NMR sample preparation for sapC-PUMA and Bcl-xL-ΔTM titration experiments

Unlabeled Bcl-xL-ΔTM was dissolved at a concentration of 900 μM at pH 6.0 for titration experiments. ^15^N SapC-PUMA (sapC-PUMA-DM) was mixed with unlabeled Bcl-xL-ΔTM to a final concentration of 100 μM of SapC-PUMA (sapC-PUMA-DM) in all experiments with increasing molar ratios of Bcl-xL-ΔTM. The following SapC-PUMA:BCL-xL-ΔTM molar ratios were tested: 1:0, 1:0.5, 1:1, and 1:2. Intensity decays of NMR signals of sapC-PUMA upon complexation were obtained with SPARKY^44^.

### Liposome fusion by Dynamic Light Scattering

Dynamic light scattering studies were conducted using a Malvern Panalytical Zetasizer Pro. Solutions of sapC at 100 μM were filtered through 50 nm pore filters and then mixed with 100 μM lipid concentration of liposomes at pH 4.2. The refractive index of liposome and sapC preparations was determined to be 1.3322 and 1.3327, respectively using an Abbe Mark III refractometer (Reichert). Absorbance values for sapC and liposomes samples obtained at the wavelength of the zetasizer laser (632.8 nm) were 0.05. The sapC and liposome mixture was immediately centrifuged for one minute at 14K rpm and the supernatant was transferred to a cuvette. The solution was allowed to equilibrate for 2 minutes before taking three measurements at each time point. Measurements were obtained every 20 minutes for a total of 200 minutes. The intensity of the scattered light at each particle size was averaged for the three measurements and plotted using QtGrace.

### Size Exclusion Chromatography

Sephacryl S-1000 superfine resin was packed in two stacked Tricorn 10/300 columns (GE Healthcare) for a final volume of 51 mL. Column assembly was equilibrated using 10 mM HEPES, 150 mM NaCl, pH 7.4. Liposomes were prepared using equilibration buffer and were injected onto the column and monitored using the following wavelengths: 240 nm, 260 nm, 280 nm, 300 nm.

### Transmission Electron Microscopy

All micrographs were obtained on a JEOL JEM 2010 transmission electron microscope with a LaB_6_ filament at 200 kV. Liposomes at 1 mM lipid concentration were prepared in 20 mM acetate buffer at pH 4.2. A total of 4 µL of these samples were deposited onto 300 mesh carbon-coated copper grids. The solution was left to dried completely or for 20 minutes and was washed three times in 40 *µ*L droplets of 1% PTA negative staining solution at pH 7. Grids were stained for 5 minutes before wiping dry. Images were taken using a Gatan camera of 1350 × 1040 pixels.

## AUTHOR INFORMATION

### Corresponding Author

*Eva de Alba. Department of Bioengineering. School of Engineering. University of California, Merced.

### Author Contributions

E.d.A., S.I.S and C.J.R wrote the manuscript. Experiments and data analysis were done by S.I.S, C.R. and E.d.A. The project was conceived by E.d.A. All authors have given approval to the final version of the manuscript.

## ACKNOWLEDGMENT

We are grateful to Mourad Sadqi for mass spectrometry measurements to check protein purity. We acknowledge Justice Lemley his help with several experiments. We are very grateful to Prof. Son Nguyen (UC Merced) for sharing with us dynamic light scattering equipment. S.I.S. and C.J.R. acknowledge graduate fellowship and undergraduate stipend support, respectively from NSF-CREST: Center for Cellular and Biomolecular Machines at the University of California, Merced (NSF-HRD-1547848).

## ABBREVIATIONS

NMR: Nuclear Magnetic Resonance
Bcl-2: B-cell lymphoma 2
Bcl-xL: B-cell lymphoma extra-large
Bax: Bcl-2 associated X
Bak: Bcl-2 antagonist killer
PUMA: p53 upregulated modulator of apoptosis
BH3: Bcl-2 homology domain 3
TEM: transmission electron microscopy
SEC: size exclusion chromatography.

## Supplementary Material

**Figure SM1.**
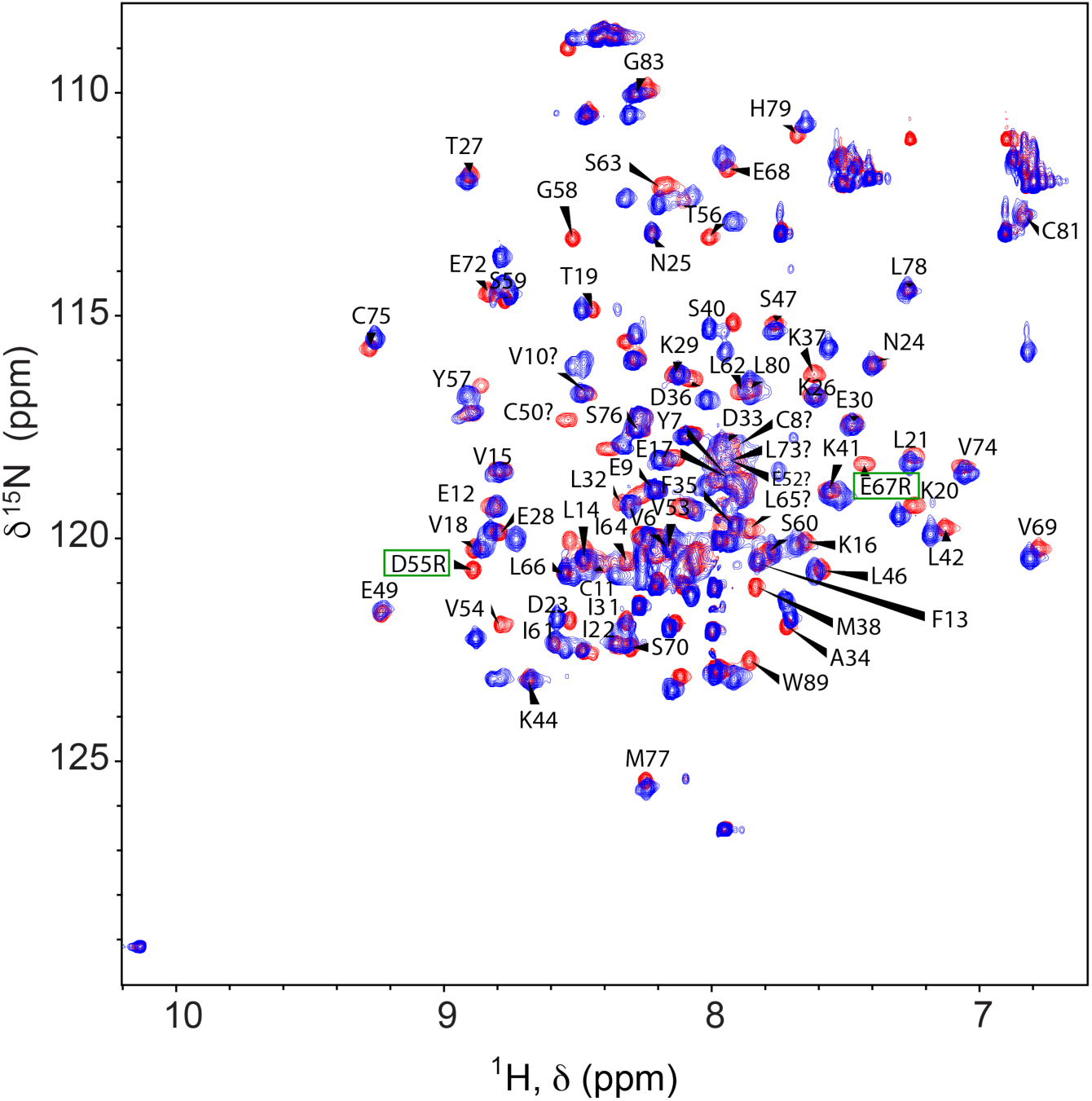
[^1^H,^15^N]-sofast HMQC of sapC-PUMA (blue) and sapC-PUMA-DM (red) at pH 6.8 The amino acids that are mutated are shown with green rectangles. Other assignments are shown with the corresponding labels.

**Figure SM2.**
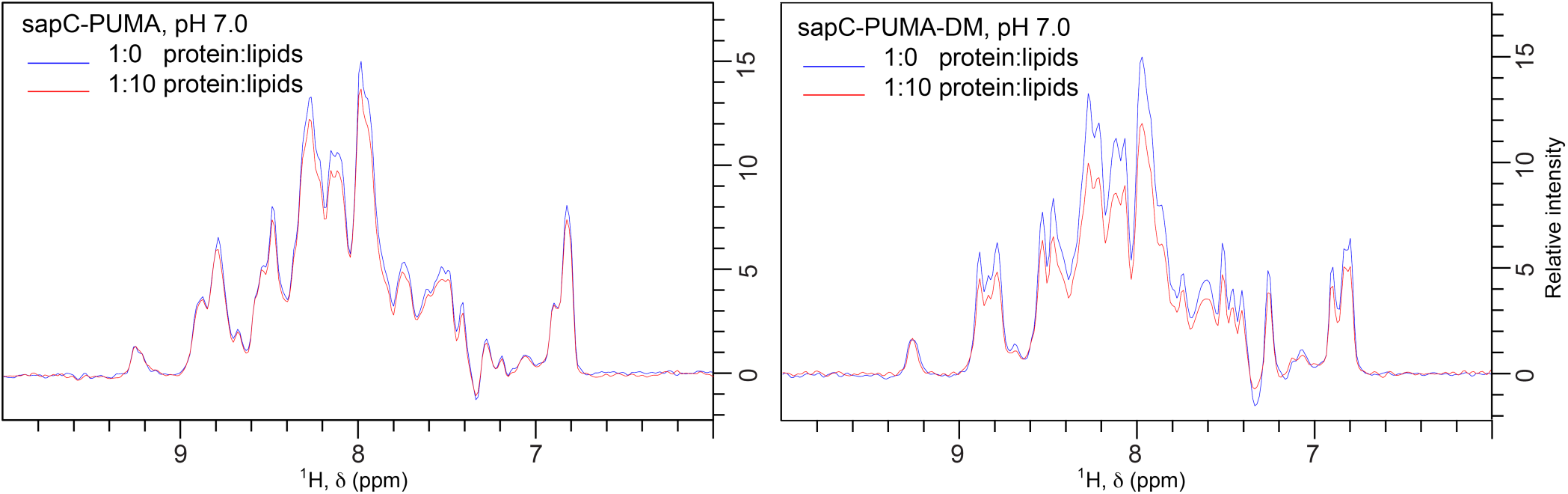
1D projections of [^1^H,^15^N]-sofast HMQC of sapC-PUMA (left panel) and sapC-PUMA-DM (right panel) in the absence (blue) and presence (red) of lipids at 1:10 protein:lipid molar ratio, pH 7. The binding of sapC-PUMA-DM is significantly larger compared to the wild-type chimera (8.5 % vs. 28 %) according to the decrease in signal to noise ratio resulting from liposome binding.

**Figure SM3.**
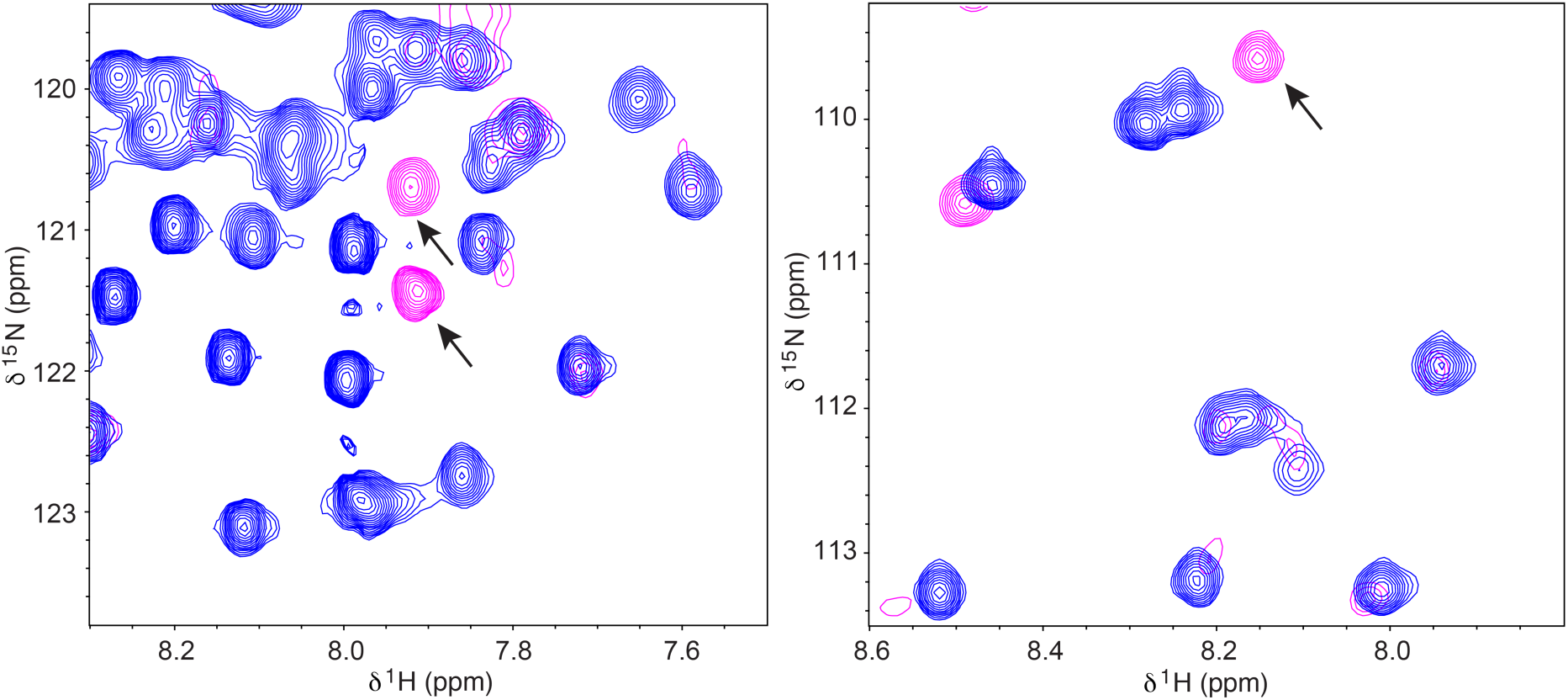
Selected regions of [^1^H,^15^N]-sofast HMQC spectra of sapC-PUMA-DM in the absence (blue) and in the presence (magenta) of 1:1 molar ratio of unlabeled Bcl-xL. The regions shown are equivalent to those appearing in Figure 8 C&D.

**Figure SM4.**
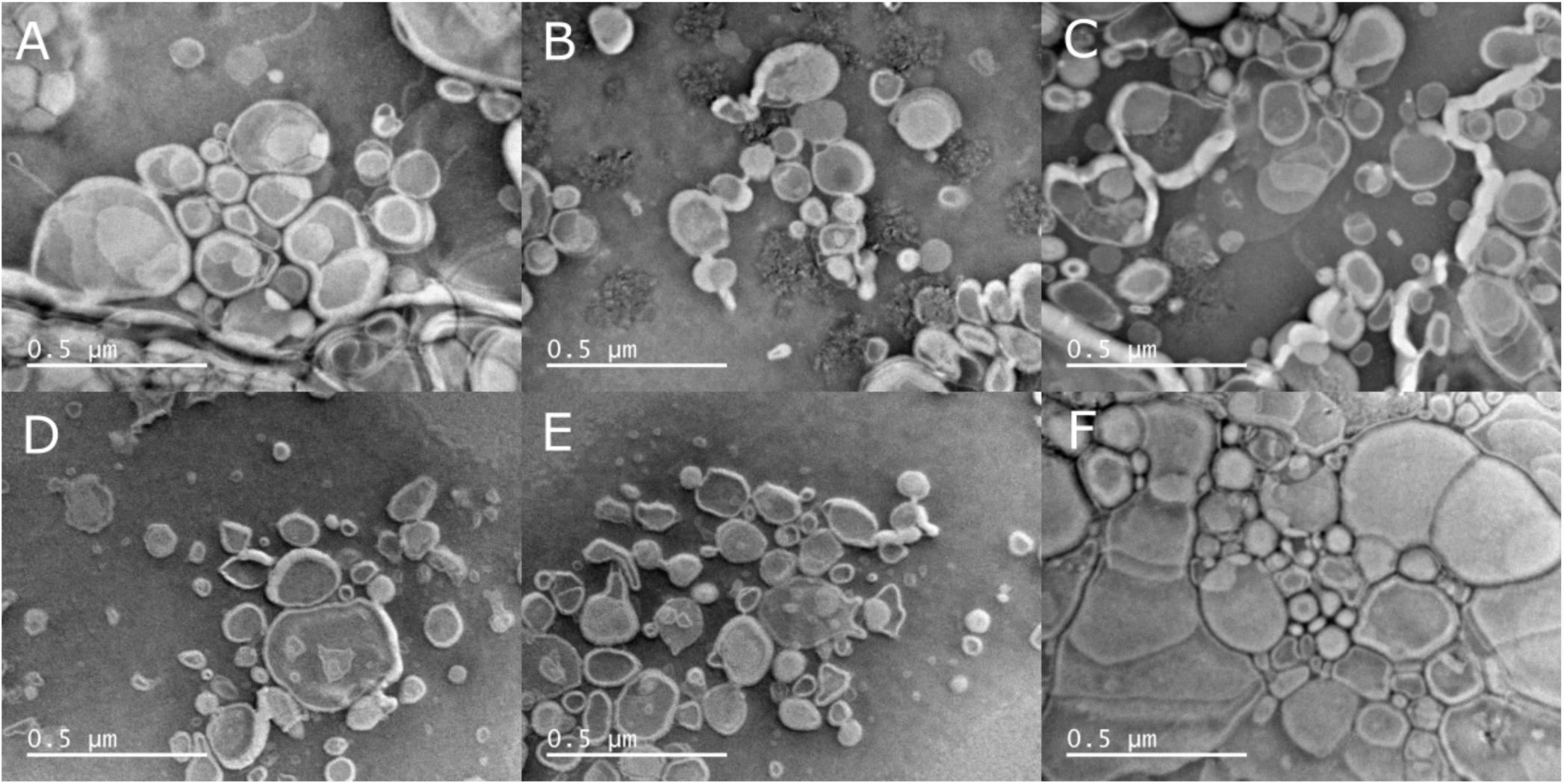
Liposome shape and size distortion in negative stained samples for TEM analysis. Micrographs A-C (same liposome sample) and F (different liposome sample but identical preparation procedure as A-C) were acquired after allowing the liposome solution to dry completely before staining. Micrographs D&E, same liposome samples as F, were allowed to dry for 20 minutes before staining.

**Figure SM5.**
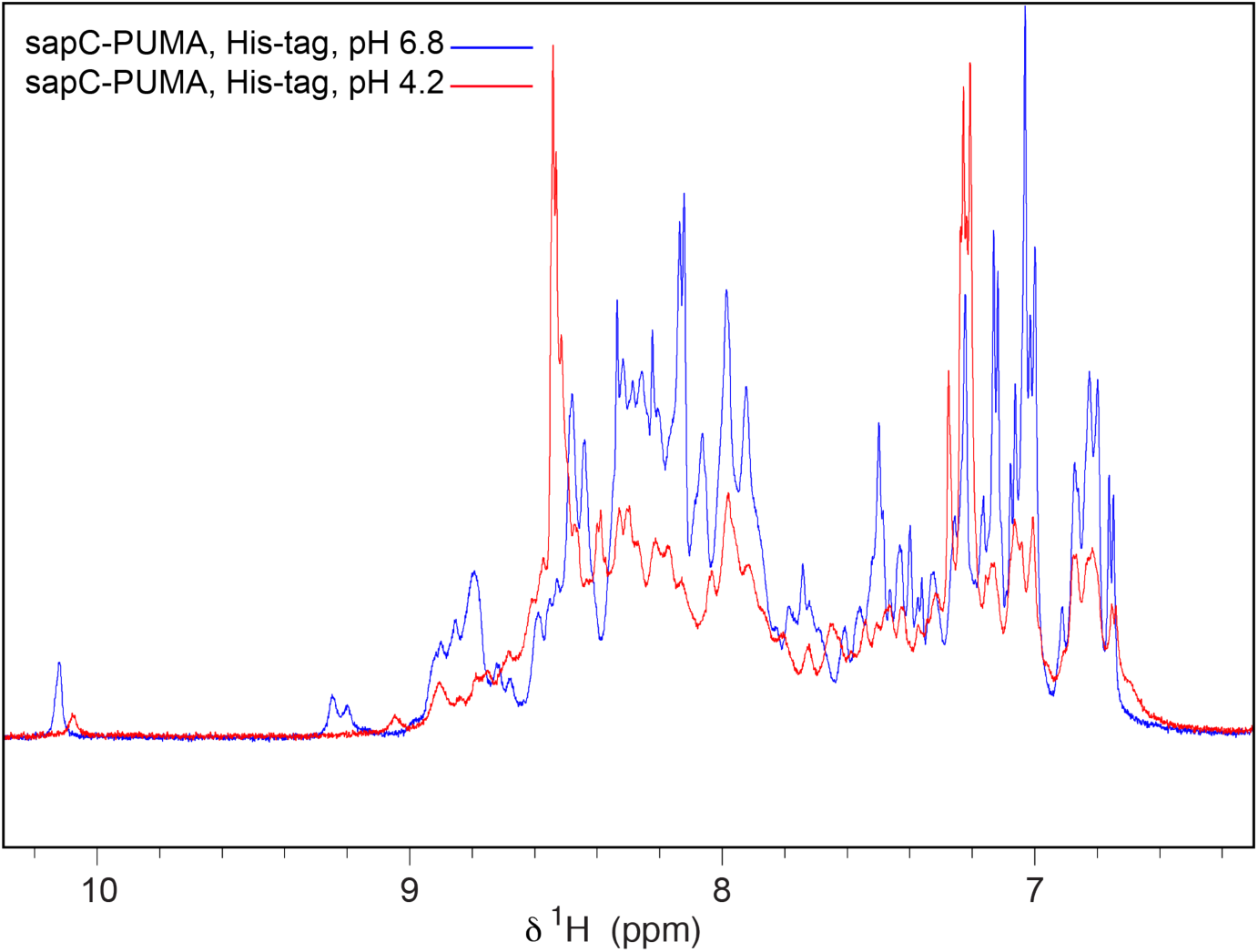
SapC-PUMA with His-tag aggregates upon acidification. SapC-PUMA with His-tag at pH 6.8 (blue, overall intensity from integration set to 100 %) and pH 4.2 (red, intensity of 77 % compared to the spectrum at pH 6.8).

